# A 1.5 Mb continuous endogenous viral region in the arbuscular mycorrhizal fungus *Rhizophagus irregularis*

**DOI:** 10.1101/2023.04.17.537115

**Authors:** Hongda Zhao, Ruixuan Zhang, Junyi Wu, Lingjie Meng, Yusuke Okazaki, Hiroyuki Hikida, Hiroyuki Ogata

## Abstract

Most fungal viruses are RNA viruses and no double-stranded DNA virus that infects fungi is known to date. A recent study detected DNA polymerase genes that originated from large dsDNA viruses in the genomes of basal fungi, suggestive of the existence of dsDNA viruses capable of infecting fungi. In this study, we searched for viral infection signatures in chromosome-level genome assemblies of the arbuscular mycorrhizal fungus *Rhizophagus irregularis*. We identified a continuous 1.5 Mb putative viral region on a chromosome in *R. irregularis* strain 4401. Phylogenetic analyses revealed that the viral region is related to viruses in the family *Asfarviridae* of the phylum *Nucleocytoviricota*. Single-copy marker genes from *Nucleocytoviricota* were detected as single-copy genes in the viral region. Furthermore, this viral region was absent in the genomes of four other *R. irregularis* strains and had fewer signals of fungal transposable elements than the other genomic regions. These results suggest a recent and single insertion of a large dsDNA viral genome in the genome of this fungal strain, providing strong evidence of the recent infection of the fungus by a dsDNA virus.

## Introduction

The fungal virosphere is dominated by RNA viruses, and a few single-stranded (ss) DNA viruses have been identified in phytopathogenic fungi (Li et al., 2020; Yu et al., 2010). Fungal viruses (i.e., mycoviruses) have been classified into 23 viral families, among which the 22 RNA virus families consist of 204 mycovirus species, while the ssDNA virus family comprises two mycovirus species (Kondo et al., 2022). No double-stranded (ds) DNA virus has been identified in fungi, but recent studies suggest they exist. A single-virion sequencing study recovered the genomes of dsDNA viruses belonging to the phylum *Nucleocytoviricota* (nucleocytoviruses) from subsurface oceanic crustal fluids, in which *Ascomycota* fungi are the main eukaryotes (Bhattacharjee et al., 2023). These viral genomes had genes that originated in fungi, suggesting that the viruses infect fungi. Additionally, DNA polymerase genes that likely originated in *Nucleocytoviricota* were also identified in the genomes of basal fungi, including *Rhizophagus irregularis*, an arbuscular mycorrhizal fungus (Gong et al., 2020). *Nucleocytoviricota* is a phylum of viruses with large dsDNA genomes (70 kb to 2.5 Mb) (Aylward et al., 2021). Their hosts include diverse eukaryotes, from protists to animals (Schulz et al., 2022).

Endogenous viral elements (EVEs) are a form or trace of viral genomes integrated into the host genome (Feschotte and Gilbert, 2012). Some previous studies identified genes from nucleocytoviruses in the genomes of eukaryotes (Maumus et al., 2014; Maumus and Blanc, 2016; Yoshikawa et al., 2019). A recent study detected giant EVEs (GEVEs) from nucleocytoviruses in the genomes of green algae (Moniruzzaman et al., 2022, 2020). A GEVE may consist of several hundred kilobases or more than a thousand kilobases, although it is often scattered across multiple contigs. The integration of nucleocytoviral genomic sequences into host genomes has significant implications for the evolution of eukaryotes, in which up to 10% of the open reading frames (ORFs) may have originated from GEVEs (Moniruzzaman et al., 2020). Such evolutionary events may be associated with the horizontal gene transfer among eukaryotic organisms (Cheng et al., 2021). Furthermore, multiple signals of virophages (dsDNA viral parasites of large DNA viruses) were detected in a variety of eukaryotic genomes (Bellas et al., 2023), reflecting the substantial diversity of the uncharacterized dsDNA virosphere in the eukaryotic domain. Analyses of EVEs may be useful for exploring the virosphere of eukaryotes and currently unidentified virus–host relationships. This is supported by previous reports describing *Nucleocytoviricota* genes in the genomes of plants (moss and fern) (Maumus et al., 2014) and oomycetes (Hannat et al., 2021), even though nucleocytoviruses have not been isolated from these organisms.

One of the difficulties in identifying GEVEs is the fragmentation of GEVEs due to the highly fragmented assemblies of eukaryotic genomes. The recently reported chromosome-level *R. irregularis* genome assemblies (Yildirir et al., 2022) enabled us to search for more complete viral fragments in the genome of this species. In the present study, we screened the genomes of five *R. irregularis* strains for *Nucleocytoviricota* signals and identified a 1.5 Mb GEVE region from a *Nucleocytoviricota*-like virus in a chromosome of strain 4401. This is the largest continuous GEVE region that has been identified to date. This GEVE is homologous to the genomes of *Asfarviridae*, a family belonging to *Nucleocytoviricota*. The results of this study provide evidence of dsDNA viruses in the fungal virosphere.

## Results

### Datasets of *R. irregularis*

We collected five chromosome-level *R. irregularis* genome assemblies corresponding to five different strains (Yildirir et al., 2022) (Table 1). These genomes were assembled using data generated by Illumina sequencing, Nanopore sequencing, and high-throughput chromatin conformation capture (Hi-C) sequencing. Long reads generated by Nanopore sequencing may be longer than a repeat or hypervariable region. In addition, Hi-C sequencing detects spatial proximity and was used for scaffolding. These technical advances have been useful for further characterizing highly fragmented eukaryotic genomes.

**Table 1.** Data regarding the *R. irregularis* genome

| Strain | Genome size (bp) | Number of scaffolds | Isolated place | Accession |
| --- | --- | --- | --- | --- |
| A1 | 147,088,061 | 33 | Switzerland | GCA_020716765.1 |
| B3 | 146,830,121 | 33 | Switzerland | GCA_020716685.1 |
| DAOM-197198 | 147,209,168 | 33 | Canada | GCA_020716725.1 |
| C2 | 161,924,360 | 33 | Switzerland | GCA_020716745.1 |
| 4401 | 146,854,905 | 33 | Canada | GCA_020716705.1 |

### Continuous 1.5 Mb GEVE region on a fungal chromosome

ViralRecall is a tool designed for detecting viral regions on the basis of the HMM profile of *Nucleocytoviricota* orthologous groups; the function used to identify 10 *Nucleocytoviricota* marker genes was integrated into this tool. By using ViralRecall to analyze the five *R. irregularis* genome assemblies, we identified a giant continuous viral signal as a 1,550 kb region (1,370,742–2,921,549 bp) on chromosome 8 of strain 4401 (Fig. 1a).

**Fig. 1.**
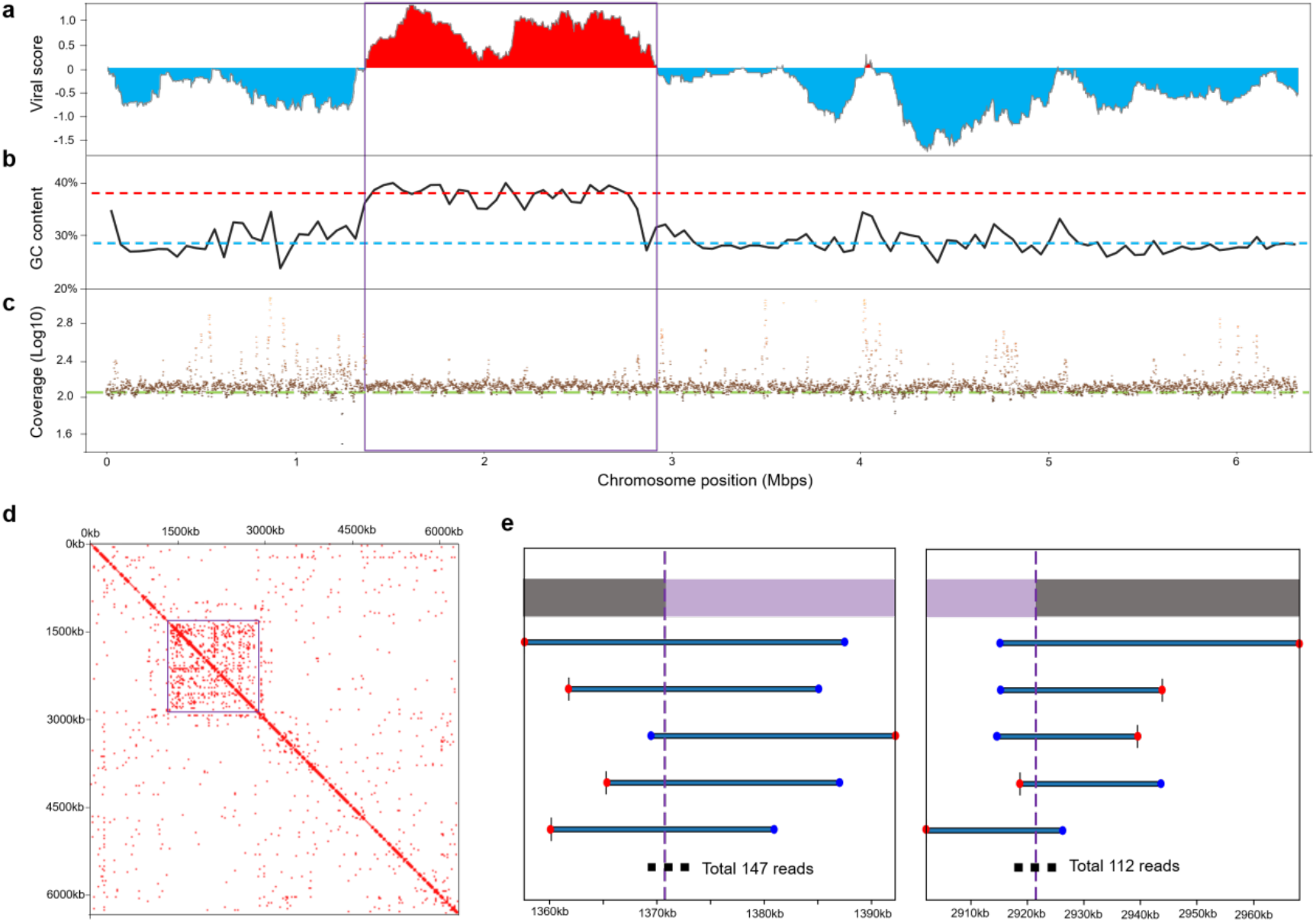
Details regarding the 1.5 Mb GEVE region of the fungal chromosome. **(a)** ViralRecall score of chromosome 8 in strain 4401. Viral scores were evaluated with a rolling window of 150 ORFs on the chromosome. The score of each ORF was based on the HMM scores of the viral reference database and cellular reference database. High and low scores represent viral and cellular regions, respectively. **(b)** GC content along the chromosome. The window size is 50,000 bp. The red and blue dashed lines represent the average GC content of the GEVE region (36.58%) and the remaining parts of chromosome 8 (28.00%), respectively. **(c)** Average read coverages of the chromosome. Each dot represents the average coverage of 1,000 continuous base pairs. The green line represents the average sequencing depth (bases/genome size = 113×). **(d)** Heat map showing Hi-C signals of chromosome 8. The resolution is 25 kb. Squares with signal values greater than 1 are marked in red and represent high contact probabilities. **(a–d)** The viral region is indicated by squares. **(e)** Examples of long reads connecting the GEVE regions identified by ViralRecall with the remaining parts of the chromosome. We present the five longest representative reads. Blue and red points indicate the start and end of the read, respectively. The extremities of the GEVE region are indicated by purple vertical dashed lines (1,370,742 bp in the figure on the left and 2,921,549 bp in the figure on the right). Purple and gray bars at the top indicate the GEVE and other cellular chromosomal regions, respectively.

This 1.5 Mb viral region had distinct sequence and structural features. Of the 10 analyzed *Nucleocytoviricota* marker genes, five were included in the 1.5 Mb viral region. These genes encode B-family DNA polymerase (PolB), RNA polymerase large subunit (RNAPL), RNA polymerase small subunit (RNAPS), mRNA capping enzyme (mRNAc), and viral late transcription factor 3 (VLTF3) (Supplementary Table S1, Supplementary Figs S1–S6). The average GC content of the *R. irregularis* genome was 27.89%, whereas the viral region had a GC content of 36.58%. The GC content throughout the region was also higher than that of the remaining parts of the chromosome (Fig. 1b). Furthermore, the Hi-C sequencing data indicated the DNA in the region was more condensed than the DNA in other genomic regions (Fig. 1d).

To confirm whether this viral region corresponds to a bona fide insertion in the fungal chromosome, we mapped the raw long reads to the fungal genome. Most chromosomal regions, including the viral region, had similar coverage (Fig. 1c). There were many short regions with a high read coverage, but they mostly corresponded to repetitive elements. There were 259 long reads directly connecting the identified viral region and cellular regions (Fig. 1e). These results confirmed that the viral region represents a GEVE integrated into the fungal chromosome.

In the GEVE region, we identified 705 ORFs, which account for 27.39% of the region. This coding density is much lower than that of other typical nucleocytoviruses. Of the 705 ORFs, 172 were functionally annotated, which revealed 53, 39, and 26 ORFs were related to “Replication, recombination, repair,” “Signal transduction mechanisms,” and “Transcription,” respectively (Supplementary Fig. S7). Using Pseudofinder, we also identified 18 putative viral pseudogenes, including the RNAPL gene, suggesting that some of the identified ORFs corresponded to pseudogenes. There were 609 intergenic regions that were longer than 100 bp (total length: 1,124 kb) (Supplementary Fig. S8). To identify traces of genes in these intergenic regions, we used these 609 intergenic sequences as queries for BLASTx searches of the NR database. Of these queries, 183 sequences (30%) matched sequences in the protein database (E-value < 10^−5^). Most of these matches (177 of the best 183 BLASTx alignments) corresponded to *Rhizophagus* protein sequences in the NR database. Moreover, more than two-thirds of these best hits were annotated as hypothetical proteins or unnamed protein products. The detection of tandem repeats in the GEVE region using Tandem Repeat Finder indicated the total length of the tandem repeats in this region was 36 kb (Supplementary Fig. S9).

### The identified GEVE is specific to strain 4401

In all five fungal strains, 59 putative viral regions were detected, including the 1.5 Mb GEVE region (Supplementary Table S2). However, with the exception of the 1.5 Mb GEVE region, all of these regions were shorter than 300 kb and contained only PolB genes (15–33 copies per genome) and/or mRNAc genes (one copy in strains A1 and B3) (Supplementary Table S3). We compared chromosome 8 in the five *R. irregularis* strains in terms of the similarity in the translated sequences (Fig. 2). Most of the strain 4401 chromosome 8 translated sequences were highly similar to the corresponding sequences in the other strains (greater than 90% sequence identity). However, the other strains lacked the region corresponding to the 1.5 Mb GEVE region in strain 4401. Consistent with this finding, chromosome 8 in strain 4401 was revealed to be approximately 1 Mb longer than chromosome 8 in the other strains (Table 2). These results indicate that the integration of the GEVE in chromosome 8 is unique to strain 4401.

**Fig. 2.**
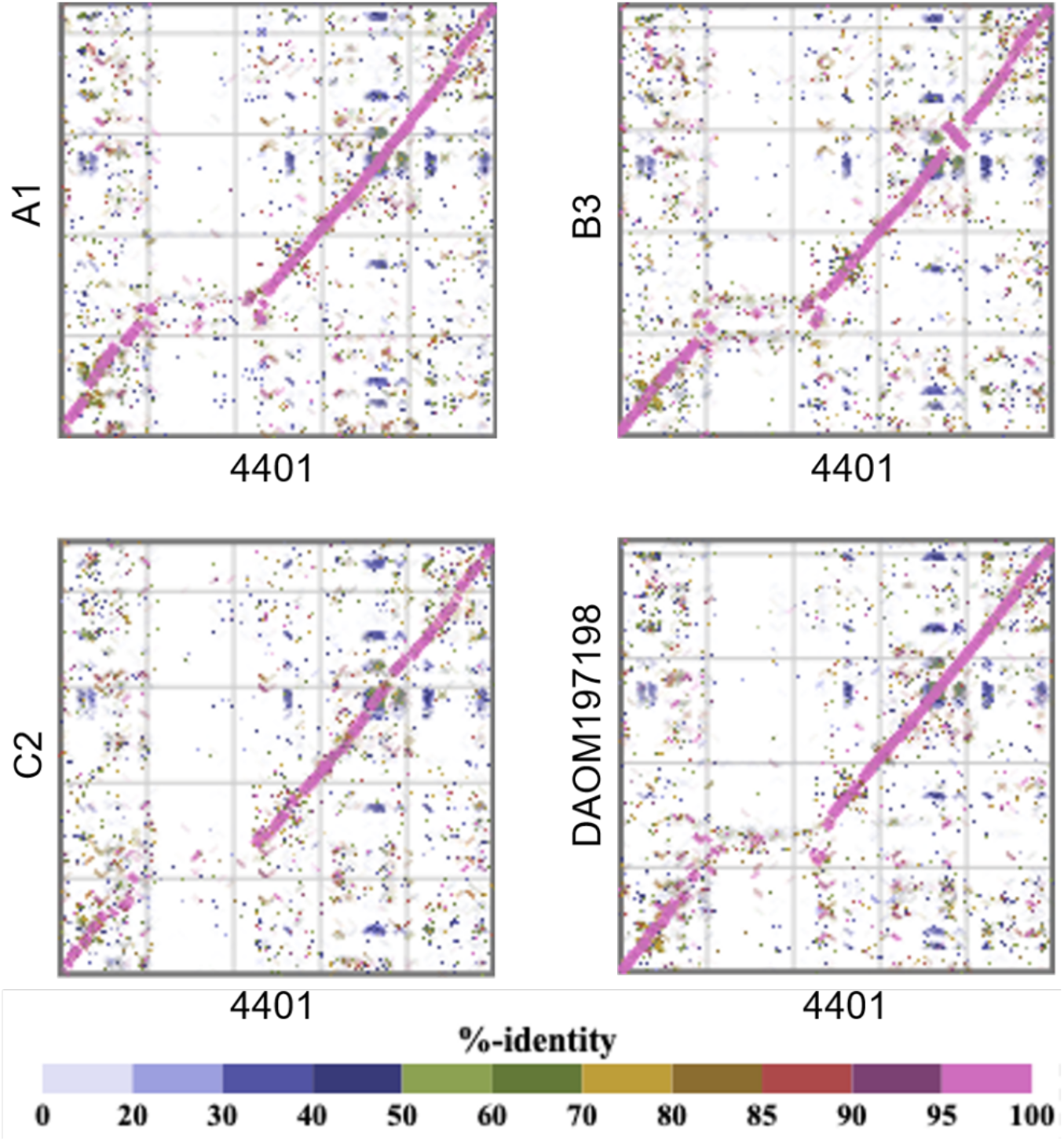
Comparison of chromosome 8 among strains. Each dot represents an amino acid sequence similarity according to tBLASTx.

**Table 2.** Chromosome 8 length in each strain

| Strain | Length (bp) |
| --- | --- |
| A1 | 5,441,258 |
| B3 | 5,350,159 |
| C2 | 5,721,097 |
| DAOM-197198 | 5,236,836 |
| 4401 | 6,327,528 |

Repeating sequences represent approximately 50% (on average) of the genomes of *R. irregularis* strains according to RepeatMasker (Yildirir et al., 2022). In the current study, we determined that 42%–65% of the individual chromosomes of the five *R. irregularis* strains consisted of repeats. In contrast, repeats represented only 18.28% of the 1.5 Mb GEVE region (5.37% in the coding region and 12.91% in the non-coding region) (Fig. 3a). The repeats in the non-coding regions accounted for 17.78% of the non-coding regions. An analysis of the repeats related to known transposable elements (TEs) indicated the TE content of the GEVE region (2.31%) was lower than that of the whole genome (average of 13.45%) (Fig. 3b).

**Fig. 3.**
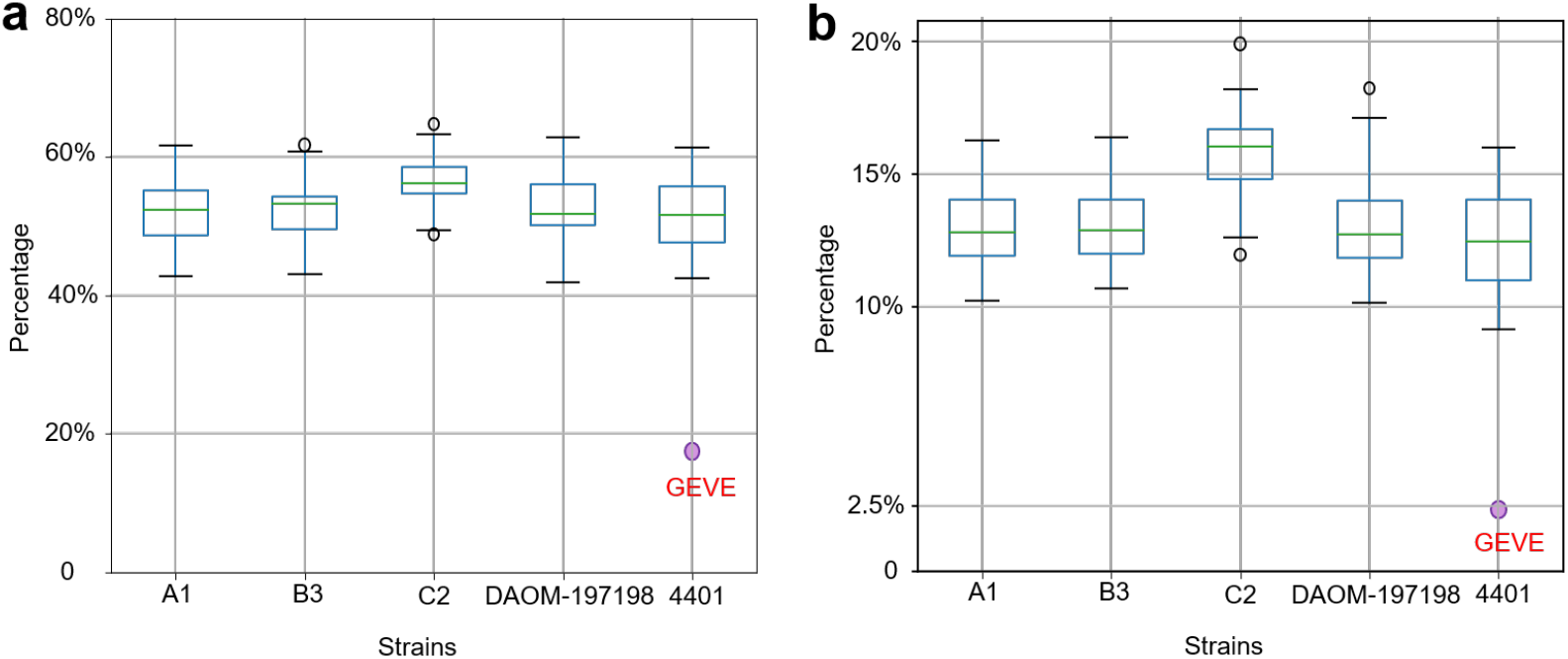
Proportion of the *R. irregularis* chromosomes comprising repeats and TEs. **(a)** Percentage of the chromosomes of the five strains and the 1.5 Mb GEVE region containing repeating sequences. The purple dot represents the repeat content in the GEVE region (18.28%). **(b)** Percentage of the chromosomes of the five strains and the 1.5 Mb GEVE region containing TE sequences. The purple dot represents the TE content in the GEVE region (2.31%). These known TEs represent a subset of the identified repeats in (a).

### The GEVE is highly homologous to *Asfarviridae* sequences

The five *Nucleocytoviricota* marker genes in the 1.5 Mb GEVE region were detected as single copies and were dispersed in the GEVE region (Fig. 4a). Phylogenetic analyses involving the five marker genes indicated that they are closely related to homologs from *Asfarviridae* (Supplementary Figs S1–S5). We performed another phylogenetic analysis using a concatenated sequence of the three longest and most universal NCLDV marker genes (PolB, RNAPL, and RNAPS) (Fig. 4b). In the constructed tree, the GEVE was classified as a sister group of the clade including *African swine fever virus* and *Abalone asfa-like virus* (100% ultrafast bootstrap support).

**Fig. 4.**
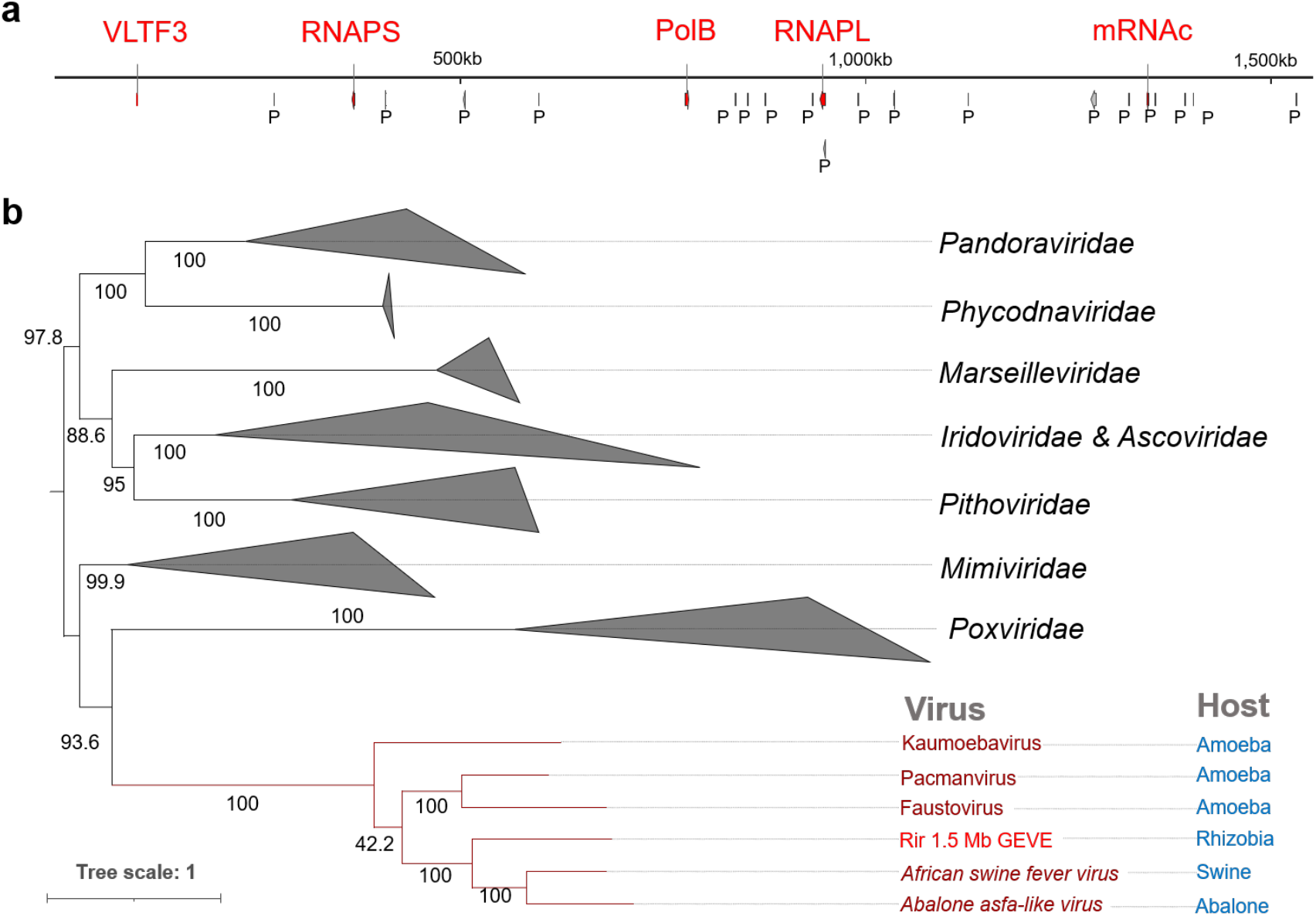
Marker genes in the 1.5 Mb GEVE region. **(a)** Distribution of *Nucleocytoviricota* marker genes and putative viral pseudogenes in the 1.5 Mb GEVE region. Marker genes are in red and viral pseudogene candidates are marked as “P”. The RNAPL gene is a marker gene and a putative pseudogene. **(b)** Concatenated maximum-likelihood phylogenetic tree constructed using three marker genes (PolB, RNAPL, and RNAPS). Rir, *R. irregularis*. Because of long branch attraction, we manually pruned the clade *Mininucleoviridae*. The root of the tree was arbitrarily chosen and the tree should be considered as an unrooted tree. Ultrafast bootstrap support values are provided along the branches. The best-fit model was Q.pfam+F+I+I+R8.

### Lack of MCP homologs in the fungal genomes

Of the 10 analyzed marker genes, the 1.5 Mb GEVE region did not include the genes encoding major capsid protein (MCP), A32-ATPase (A32), D5 primase/helicase (D5), ribonucleotide reductase (RNR), and superfamily II helicase (SFII). These five genes were not detected in the genomes of the five strains. We found homologs of the *Asfarviridae* minor capsid protein-encoding gene in the GEVE region (171,954–176,120 bp). We performed tBLASTn and other searches using HMM models of MCP sequences from multiple viral groups, including typical MCPs of nucleocytoviruses (see Methods for details). However, MCP genes were not detected in strain 4401 or the other four strains.

### The fungal GEVE is closely related to virus-like sequences from a sea slug

We performed a BLASTp search of the NR database using the 705 predicted ORFs in the 1.5 Mb GEVE region as queries. On the basis of the best matches, we determined the most likely taxonomic distribution of 184 ORFs (E-value < 10^−5^). More specifically, 102 ORFs were most similar to eukaryotic sequences, 56 ORFs were most similar to prokaryotic sequences, and 26 ORFs were most similar to viral sequences (21 of the viral sequences were from *Asfarviridae*). We manually checked the ORFs that closely matched prokaryotic sequences. There were 21 ORFs that were most similar to bacterial genes, but were also similar to *Asfarviridae* genes. Because the prokaryotic sequences that were most similar to these ORFs were environmental sequences, we speculated that these ORFs are viral ORFs (Fig. 5a). A similar taxonomic distribution was observed for previously reported nucleocytoviruses lacking close relatives in databases (Blanc-Mathieu et al., 2021).

**Fig. 5.**
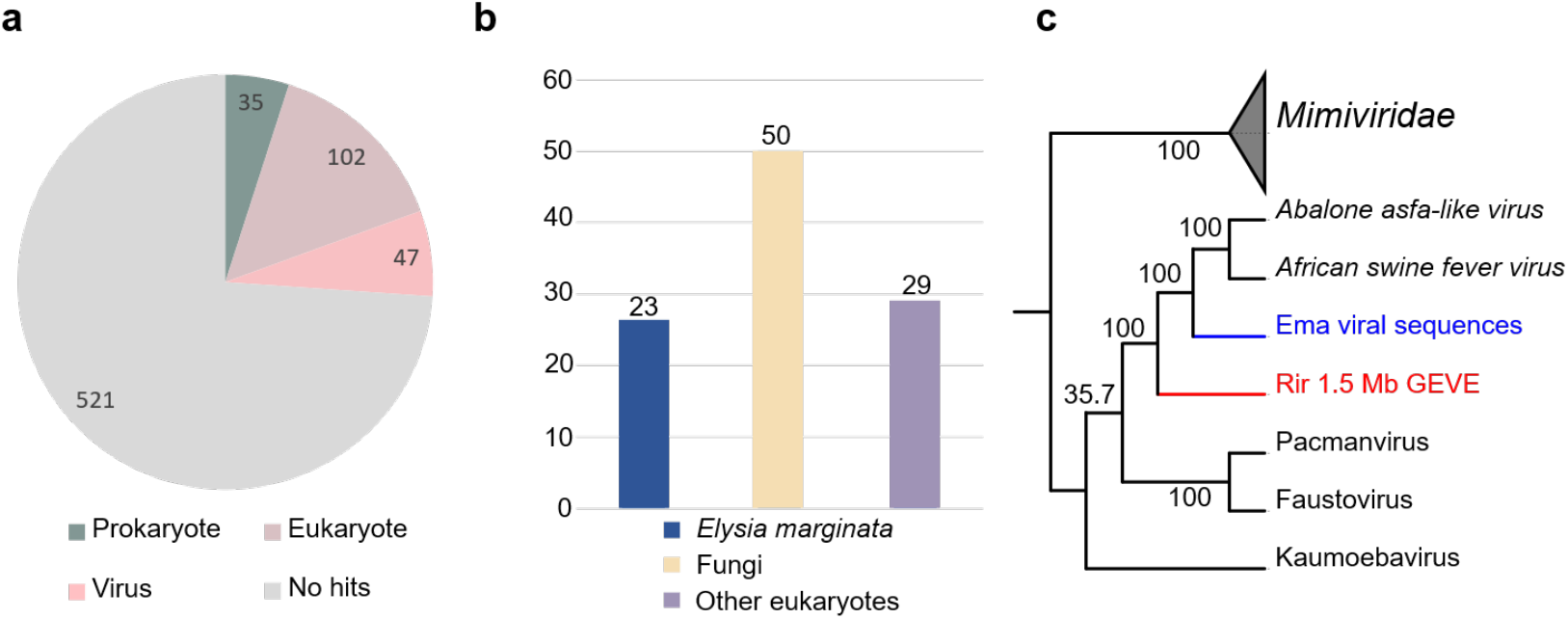
Relationship between the GEVE on chromosome 8 of strain 4401 and viral sequences in *E. marginata*. **(a)** Taxonomic distribution of the 705 ORFs in the 1.5 Mb GEVE region according to the best matches revealed by the BLASTp search of the edited NR database. Twenty-one ORFs that most closely matched prokaryotic genes were designated as viral genes (see main text for details). **(b)** Details regarding 102 eukaryotic annotations. The 23 best matches to *E. marginata* sequences were from one assembly (GCA_019649035.1). **(c)** Maximum-likelihood phylogenetic tree of the viral sequences from the *E. marginata* genome constructed using three concatenated marker genes (PolB, RNAPL, and RNAPS). *Mimiviridae* sequences were selected as the outgroup. Rir, *R. irregularis*; Ema, *E. marginata*. The best-fit model was Q.pfam+F+I+G4.

Notably, 23 of the 102 ORFs that most closely matched eukaryotic sequences were most similar to a single genome assembly of a sea slug species (*Elysia marginata*) (Maeda et al., 2021) (Fig. 5b). However, most of the following matches for these ORFs were genes from viruses belonging to *Asfarviridae*, suggestive of the presence of viral-like sequences in the *E. marginata* genome. By screening the *E. marginata* genomic data, we detected 13 putative viral regions and 10 *Nucleocytoviricota* marker genes (PolB, RNAPS, RNAPL, MCP, RNR, VLTF3, A32, SFII, D5, and mRNAc) (Supplementary Tables S1 and S4). Six regions had a GC content of approximately 35%, which was similar to the GC content of *E. marginata* (36.5%), but seven regions had a higher GC content (56.7%). The total length of these seven regions was 124 kb and included all of the identified marker genes. The length of each region closely matched the length of the corresponding contig.

A phylogenetic analysis of the marker genes confirmed that the viral sequences in *E. marginata* are related to *Asfarviridae* and the GEVE in *R. irregularis* (Fig. 5c). In the phylogenetic tree, the viral sequences from *E. marginata* were positioned between the fungal GEVE and the clade including *African swine fever virus* and *Abalone asfa-like virus*. Consistent phylogenetic relationships were also detected among each marker gene (Supplementary Figs S1– S6). Thus, the *E. marginata* genomic data contains sequences related to *Asfarviridae*.

## Discussion

The fungal virosphere, which mostly comprises RNA viruses, lacks dsDNA viruses (Kondo et al., 2022). In the present study, we identified a continuous 1.5 Mb *Nucleocytoviricota* GEVE region in the arbuscular mycorrhizal fungus *R. irregularis*. Prior to this study, the longest known GEVE (475 kb) was detected in a single genomic contig from a green alga (Moniruzzaman et al., 2022). Thus, this GEVE region in a fungal chromosome represents the longest continuous GEVE in a eukaryotic genome identified to date. Our phylogenetic analysis indicated this GEVE is closely related to the clade containing *African swine fever virus* and *Abalone asfa-like virus* (Fig. 4b, Supplementary Figs S1–S5). Additionally, the GEVE region includes single copies of five *Nucleocytoviricota* marker genes (Fig. 4a) and its GC content and Hi-C profile differ from those of the other parts of the chromosome (Fig. 1b and d). These results suggest that the GEVE region originated from a single insertion of an *Asfarviridae*-related virus. The compaction of the GEVE genomic region revealed by the Hi-C data may reflect the silencing of genes in this region, which was previously observed in mammals (Shukron et al., 2019). Such genomic compaction may function as a defense of fungi against the invasion of extracellular DNA. The GEVE was detected in only one of the five analyzed strains (Fig. 2, Supplementary Table S2). This GEVE region had a lower repeat and TE density than the other fungal chromosomal regions, implying that hat the region experienced a shorter time for invasions of TEs than the other chromosomal regions (Fig. 3). This viral integration was probably a relatively recent event, suggesting there are dsDNA viruses that are still actively infecting the fungal species.

The genomes of isolated *Asfarviridae* viruses (155–466 kb) (Matsuyama et al., 2020; Reteno et al., 2015) are much smaller than the 1.5 Mb GEVE region identified in the fungal chromosome. Furthermore, the coding density of the GEVE region was low (27.39%). The expansion of repeated sequences after a viral integration may help to explain the expansion of the GEVE region, but it does not appear to fully account for the high proportion of non-coding sequences in the GEVE (72.61%) because repeats represented 17.78% of the non-coding regions, which also contained some tandem repeats. The substantial abundance of non-coding sequences in the GEVE is probably related to the decay of coding sequences, with one-third of the intergenic sequences containing traces of genes (Supplementary Fig. S9).

Although the long and contiguous GEVE region was detected only in chromosome 8 of strain 4401, PolB genes that may have originated in *Asfarviridae* were identified on multiple chromosomes in all strains (Supplementary Table S1). The genomic positions and copy numbers of these PolB genes varied among the genomes of the analyzed *R. irregularis* strains (Supplementary Table S3). This suggests that infections of *R. irregularis* by *Asfarviridae*-like viruses may be a widespread and on-going event.

*Asfarviridae* viruses infect a variety of eukaryotes, such as swine, abalone, and amoebae, although only five have been isolated and completely sequenced (Karki et al., 2021). Previous studies identified marine dinoflagellates (Ogata et al., 2009) and oomycetes (Hannat et al., 2021) as potential hosts of *Asfarviridae*. In the present study, we detected *Asfarviridae*-like genome sequences closely related to the fungal GEVE within the *E. marginata* genome (Fig. 4c). The viral sequences in *E. marginata* encode all of the core genes (Supplementary Table S1), suggesting that the virus containing these sequences can form virion particles and is infectious. Unlike the GEVE in *R. irregularis*, the seven viral regions with high GC contents in *E. marginata* covered almost the entire length of their contigs, suggesting that these viral regions are not insertions in the sea slug genome, but were derived from viral genomes concomitantly sequenced with the *E. marginata* genome (Supplementary Table S2). Considered together, these findings indicate *Asfarviridae* viruses are likely more widespread and diverse than previously believed.

Recent research confirmed MCP is one of the major components of virions (Krupovic et al., 2022), but an MCP gene was not detected in the *R. irregularis* genome (Supplementary Table S1). However, we identified *Asfarviridae* minor capsid protein-encoding homologs in the 1.5 Mb GEVE region, implying this virus may have a capsid structure. There are two possible explanations for the lack of MCP genes. First, some genomic regions may have been deleted or genomic rearrangements might have occurred after the viral genome was integrated (Moniruzzaman et al., 2020), resulting in a lack of an MCP gene in the GEVE region. However, this is unlikely because the GEVE is nearly complete (with many conserved single-copy marker genes) and MCP homologs were not detected in the genomes of the five analyzed strains. Second, the viral genome from which the GEVE was derived may include an MCP gene whose sequence differs from that of the known MCP genes. The MCPs of nucleocytoviruses are highly diverse and some nucleocytoviruses (e.g., pandoraviruses) use different types of proteins as the major component of the virion (Krupovic et al., 2020). Accordingly, unknown MCP genes may not have been detected in the current study because our detection methods were based on known reference sequences.

Arbuscular mycorrhizal fungi are obligate plant mutualistic organisms that provide significant benefits to plants (e.g., increased nutrient levels and enhanced disease resistance) (Gosling et al., 2006). As a model organism of arbuscular mycorrhizal fungi, *R. irregularis* forms a robust tripartite association with its endobacteria and plants during its life cycle; horizontal gene transfers among these organisms contribute to their evolution and symbiotic adaptation (Li et al., 2018). For example, foreign genes affect *R. irregularis* life cycle-related processes, including gene expression, mitosis, and signal transduction (Li et al., 2018). In the present study, we revealed that dsDNA viruses may also be important for the horizontal transfer of genes in *R. irregularis*. Future functional analyses of these virus-derived genes in *R. irregularis* may provide novel insights into the ecology and evolution of this beneficial microorganism.

## Methods

### Detection of viral regions

To identify virus-like regions in eukaryotic genomes, we used ViralRecall v2.1 (Aylward and Moniruzzaman, 2021) to screen genomic data, with a window size and viral score set at 150 ORFs and 0, respectively. ViralRecall is a tool designed for identifying virus-like regions. Notably, this tool uses *Nucleocytoviricota* orthologous groups and the Pfam database to detect *Nucleocytoviricota* signals. This tool evaluates the viral and cellular scores for each ORF in the genome and determines the likelihood these regions are virus-like. Chromosome-level genomic data for five *R. irregularis* strains were retrieved from GenBank, whereas *E. marginata* genomic data were retrieved from GCA_019649035.1 (Maeda et al., 2021). The GC content was calculated using in-house Python scripts, with the window size set at 50,000 bp to minimize the effect of repeats. The insertion of the viral region in the fungal chromosome was validated as follows. First, long reads of strain 4401 (accession no. SRR15461860) were mapped to the whole genome using Minimap2 v2.24 (Li, 2018). We applied the “view” function of Samtools v1.16.1 (Li et al., 2009) to select reads connecting the virus and host region. The “bamtobed” function of Bedtools v2.29.2 (Quinlan and Hall, 2010) was used to visualize the results. The comparison of chromosome 8 from different strains (at the amino acid sequence level) was performed using DiGAlign v1.3 (http://www.genome.jp/digalign/).

### Generation of Hi-C contact map and detection of repeats

To clarify the structure of the chromosome with the 1.5 Mb GEVE region, we transformed the raw Hi-C sequencing data of strain 4401 from the NCBI Sequence Read Archive (accession no. SRR15461854) into normalized contact maps using Juicer v2.0 (Durand et al., 2016) and visualized the results using Juicebox (http://aidenlab.org/juicebox/). Repetitive elements in the genome were identified and masked using RepeatModeler v2.0.2 (Flynn et al., 2020) and RepeatMasker v4.1.2 (http://www.repeatmasker.org). RepeatModeler can identify repeats (in both coding and non-coding regions) and annotate TEs, including retroelements and DNA transposons, with distinct discovery algorithms. We used the “-LTRStruct” parameter while running RepeatModeler to detect long terminal repeat retroelements.

### Phylogenetic analyses

*Nucleocytoviricota* marker genes were predicted using the built-in HMMER profile of ViralRecall. To verify whether these genes are indeed viral homologs, we obtained reference sequences of these marker genes from a previous study (Kazlauskas et al., 2020) and GenBank and then constructed phylogenetic trees as follows. A multiple sequence alignment was completed using Clustal-Omega v1.2.4 (Sievers and Higgins, 2018) and trimmed using trimAl v1.4.1 (parameter: “-gt 0.1”) (Capella-Gutiérrez et al., 2009). Maximum-likelihood phylogenetic trees were generated using IQ-TREE v2.2.0 (Minh et al., 2020), with 1,000 ultrafast bootstrap replicates (Hoang et al., 2018). The best-fit model was selected using ModelFinder (Kalyaanamoorthy et al., 2017). Phylogenetic trees were visualized using iTOL v6.7.4 (Letunic and Bork, 2019).

We screened the genome of strain 4401 for MCP genes using hmm files constructed from the MCP sequences of phages, *Nucleocytoviricota, Mirusviricota*, and *Herpesvirales* as well as HMMER v3.3.2 (Eddy, 2011) (e < 0.05). We also searched for major virion capsid 1 and 2 of pandoraviruses using the same method. Reference sequences of phages, *Herpesvirales*, pandoraviruses, and other *Nucleocytoviricota* viruses were obtained from GenBank and the NCBI protein database. Previously reported MCP sequences of *Mirusviricota* were also used (Gaïa et al., 2023). Considering the possibility of missing results if a sequence in the fungal genome is not identified as an ORF, we used NCBI BLAST to perform a tBLASTn search (Johnson et al., 2008) of the *R. irregularis* nucleotide sequences (NCBI: txid588596) in the nr/nt database to identify MCP sequences (E-value < 0.05).

### Annotation of viral regions

The ORFs in the GEVE were predicted using ViralRecall in Prodigal v2.6.3 (Hyatt et al., 2010). As a tool designed for identifying prokaryotic genes, it efficiently predicted viral genes on the basis of the similarity in the genomic architecture, but it may not be ideal for predicting some eukaryotic genes (Moniruzzaman et al., 2020). The ORFs were annotated using BLASTp in BLAST+ v2.13.0 (Camacho et al., 2009) (E-value < 10^−5^). Because previous studies may have annotated viral insertions in fungal genomes as fungal genes, we used the NR database lacking the sequences from the fungal class *Glomeromycetes* (NCBI: txid214506, which includes *R. irregularis*). The best match for each ORF was used to determine the taxonomic distribution of the ORFs (i.e., eukaryote, prokaryote, and virus). Functional annotations were retrieved using eggNOG-mapper v2.1.9 (Cantalapiedra et al., 2021).

Pseudogene candidates were predicted using Prokka v1.14.6 (Seemann, 2014) and Pseudofinder v1.1.0 (Syberg-Olsen et al., 2022), with Viral-Host database 11/2022 (Mihara et al., 2016) serving as the reference. To identify traces of genes, we extracted the genomic region between two predicted ORFs and eliminated the regions shorter than 100 bp before performing a BLASTx search using Diamond v2.0.15 (E-value < 10^−5^) (Buchfink et al., 2015). The NR database was used as the reference and “--ultra-sensitive” was selected as the parameter. We also used Tandem Repeat Finder v4.09 (Benson, 1999) to identify the tandem repeats in the GEVE region, with “2 7 7 80 10 30 2000 -f -d -m” selected as the parameter.

## Acknowledgments

This work was supported by JSPS/KAKENHI (Nos. 22H00384 and 19H05667 to H.O.). Computational time was provided by the Supercomputer System, Institute for Chemical Research, Kyoto University. We thank Edanz (https://jp.edanz.com/ac) for editing a draft of this manuscript.

## Author Contributions

HZ performed all of the analyses and wrote the first draft of the manuscript. RZ and JW helped conceive this study. LM contributed to the identification of the 1.5 Mb GEVE region. YO, HH, and HO supervised HZ and helped finalize the manuscript. All authors read and approved the final version of the manuscript.

## Supplementary figures

**Supplementary Fig S1:**
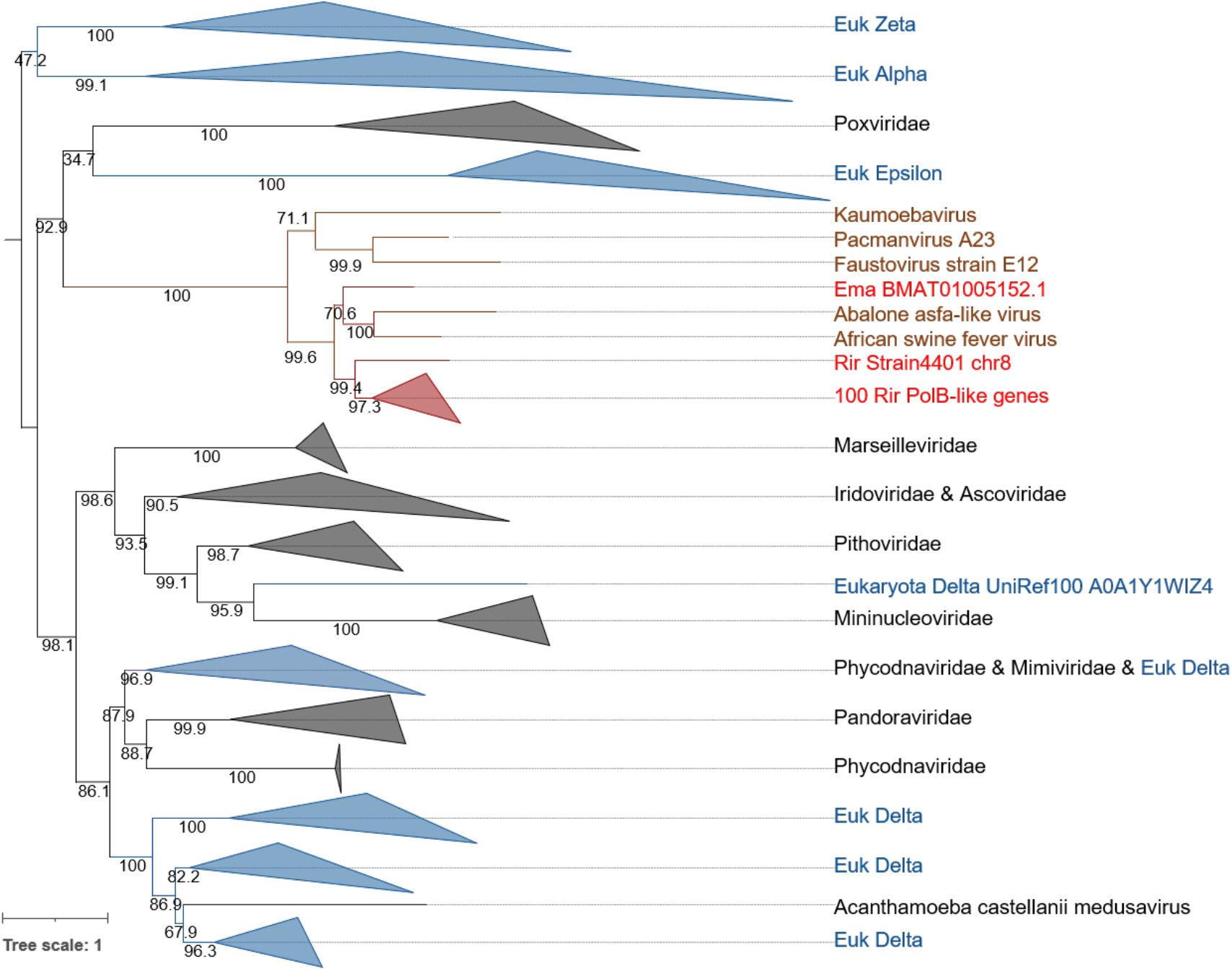
Phylogenetic tree of 101 PolB candidates predicted by ViralRecall. Clades containing eukaryotic sequences are in blue. *Asfarviridae* sequences are in brown. Viral sequences from *R. irregularis* and *E. marginata* are indicated by red labels. Ultrafast bootstrap support values are provided along the branches. These 100 PolB sequences are phylogenetically distinct from the one detected in the 1.5 Mb GEVE region. The root of the tree was arbitrarily chosen and the tree should be considered as an unrooted tree. The best-fit model was Q.pfam+F+R10.

**Supplementary Fig S2:**
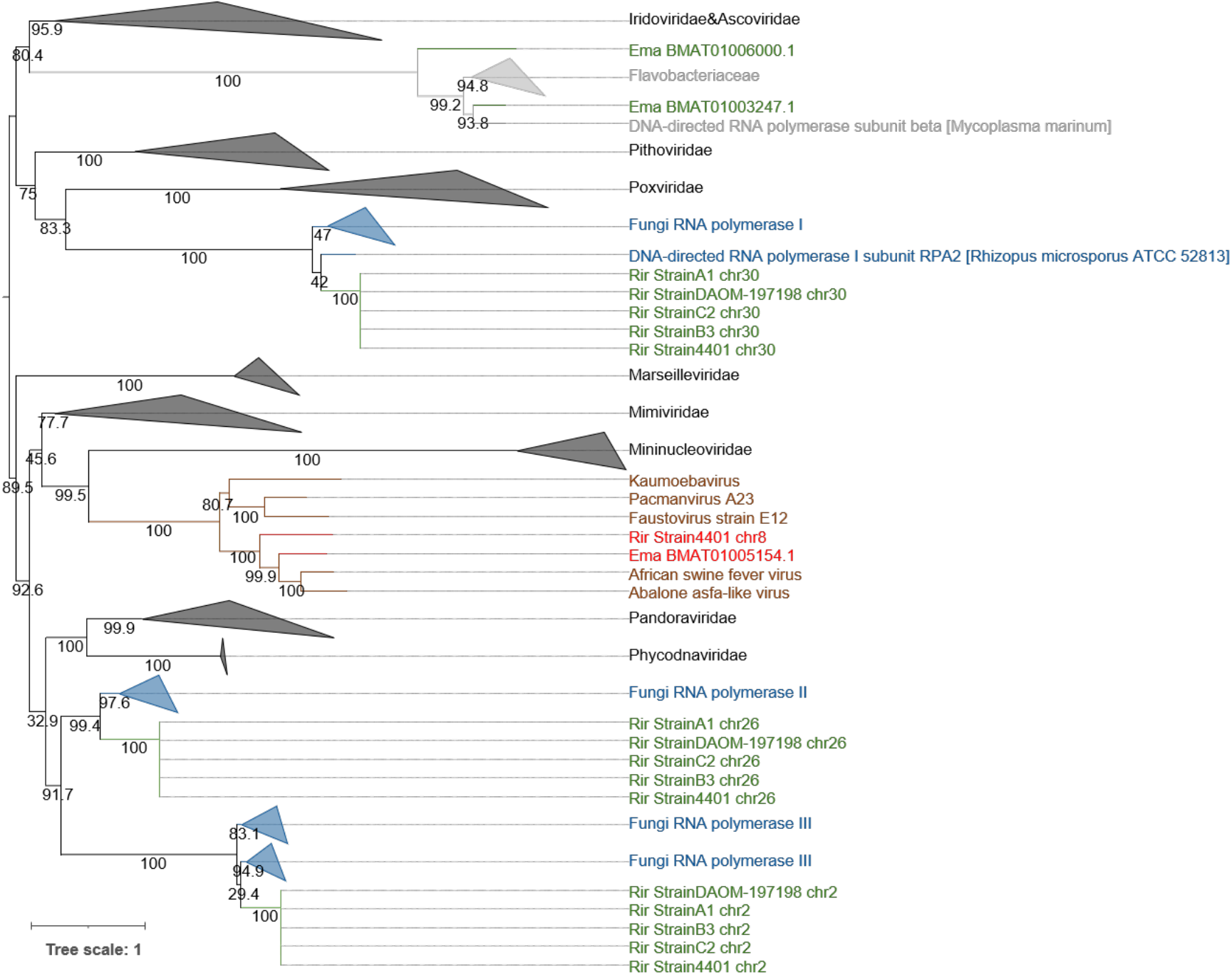
Phylogenetic tree of RNAPS candidates predicted by ViralRecall. The Fig. S1 legend explains the meaning of the different colors; however, eukaryotic and bacterial sequences in *R. irregularis* and *E. marginata* are in green and bacterial sequences are in light gray. Ultrafast bootstrap support values are provided along the branches. The root of the tree was arbitrarily chosen and the tree should be considered as an unrooted tree. The best-fit model was LG+F+I+I+R8.

**Supplementary Fig S3:**
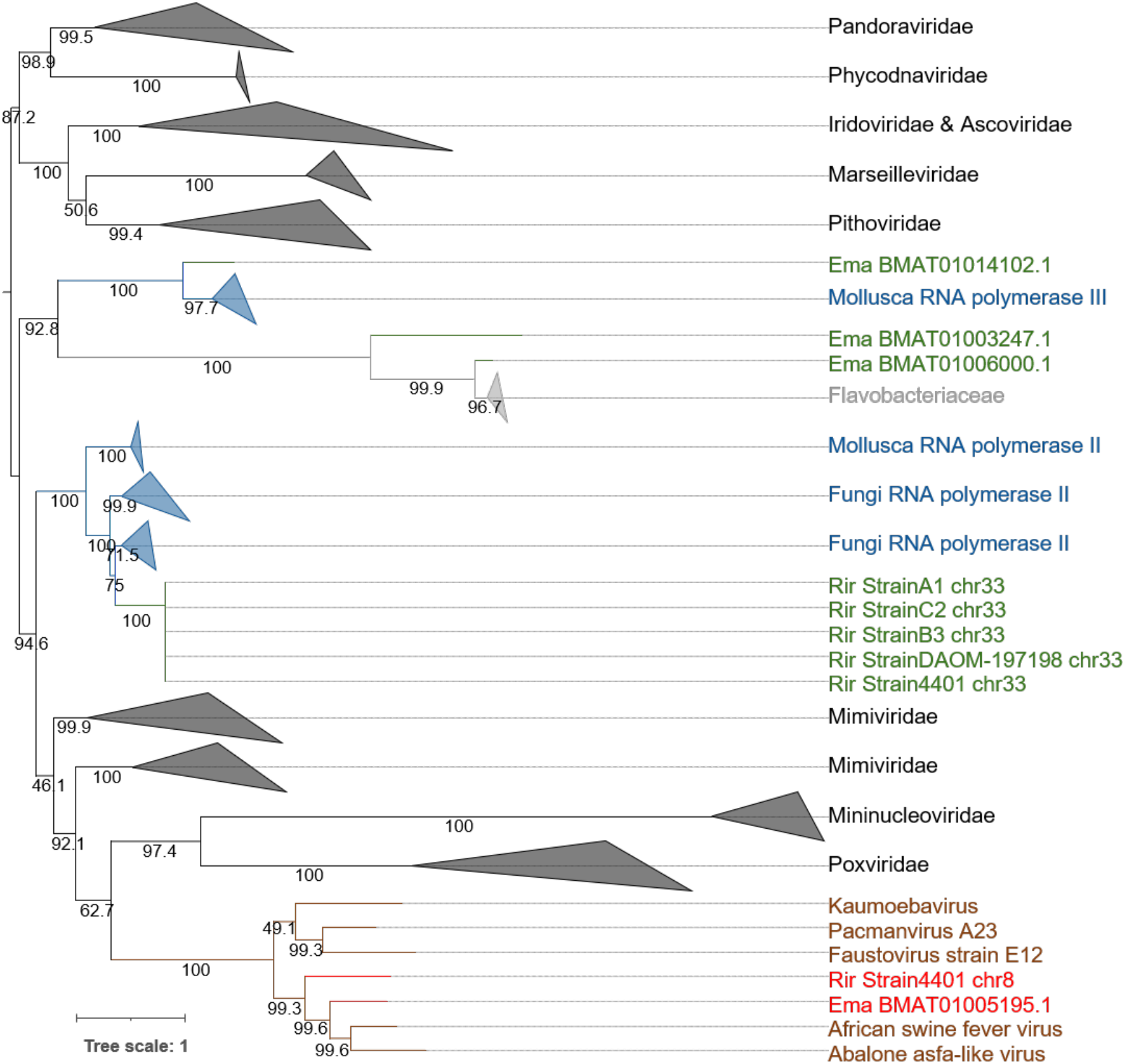
Phylogenetic tree of RNAPL candidates predicted by ViralRecall. The Fig. S2 legend explains the meaning of the different colors. Ultrafast bootstrap support values are provided along the branches. The root of the tree was arbitrarily chosen and the tree should be considered as an unrooted tree. The best-fit model was LG+F+R10.

**Supplementary Fig S4:**
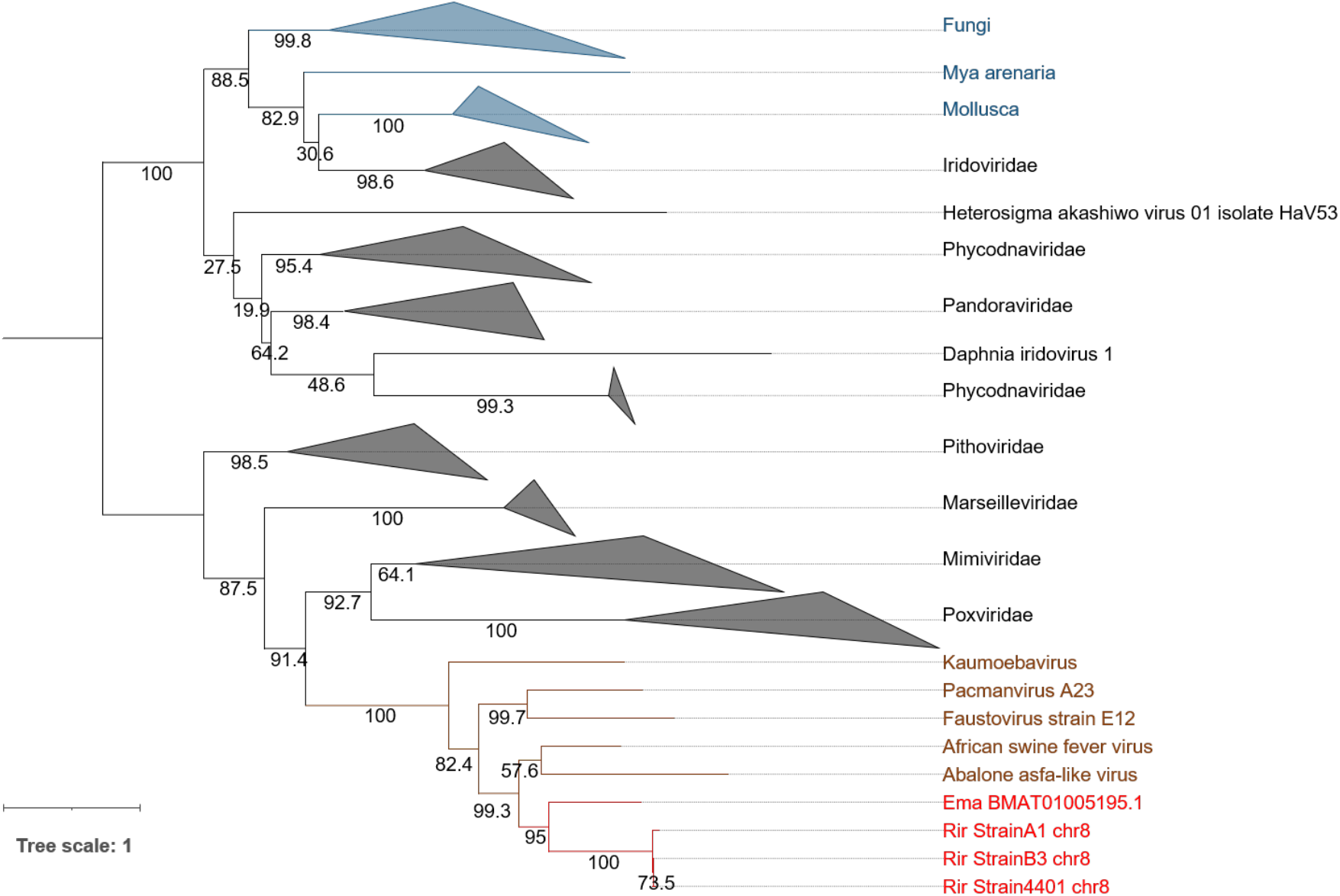
Phylogenetic tree of mRNAc candidates predicted by ViralRecall. The Fig. S2 legend explains the meaning of the different colors. Ultrafast bootstrap support values are provided along the branches. The root of the tree was arbitrarily chosen and the tree should be considered as an unrooted tree. The best-fit model was Q.pfam+F+I+I+R6.

**Supplementary Fig S5:**
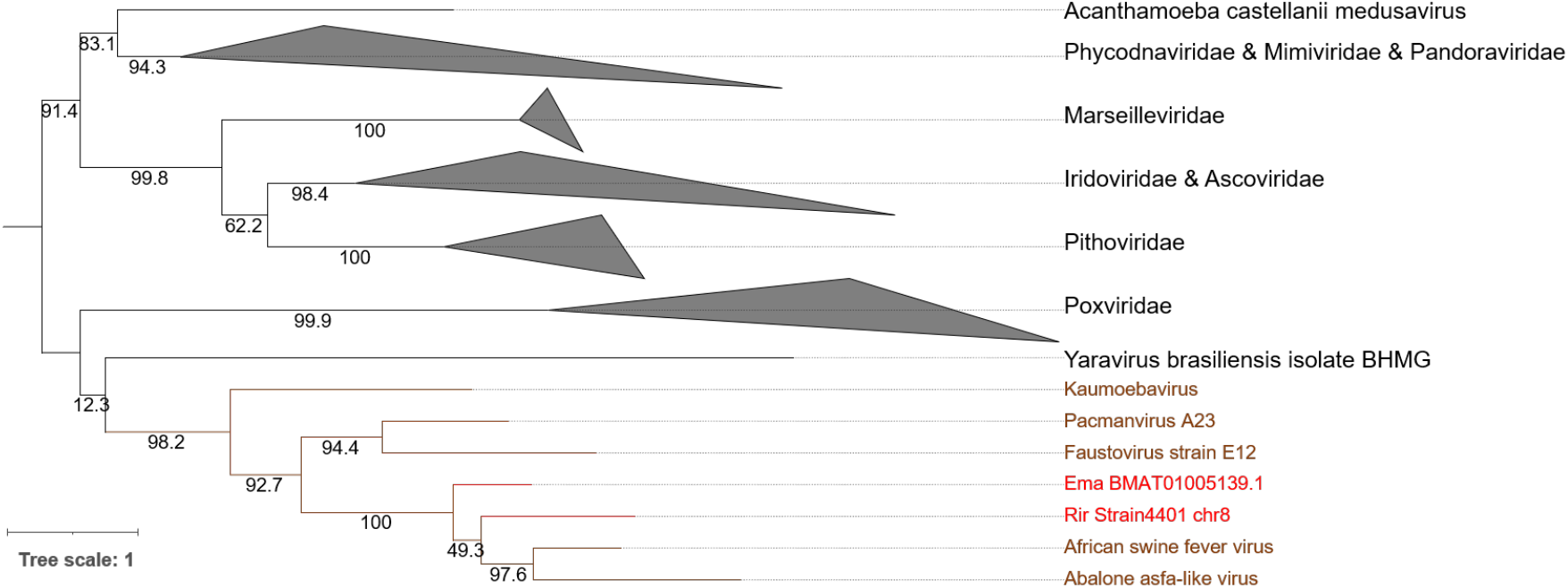
Phylogenetic tree of VLTF3 candidates predicted by ViralRecall. The Fig. S1 legend explains the meaning of the different colors. Ultrafast bootstrap support values are provided along the branches. The root of the tree was arbitrarily chosen and the tree should be considered as an unrooted tree. The best-fit model was Q.insect+F+R6.

**Supplementary Fig S6:**
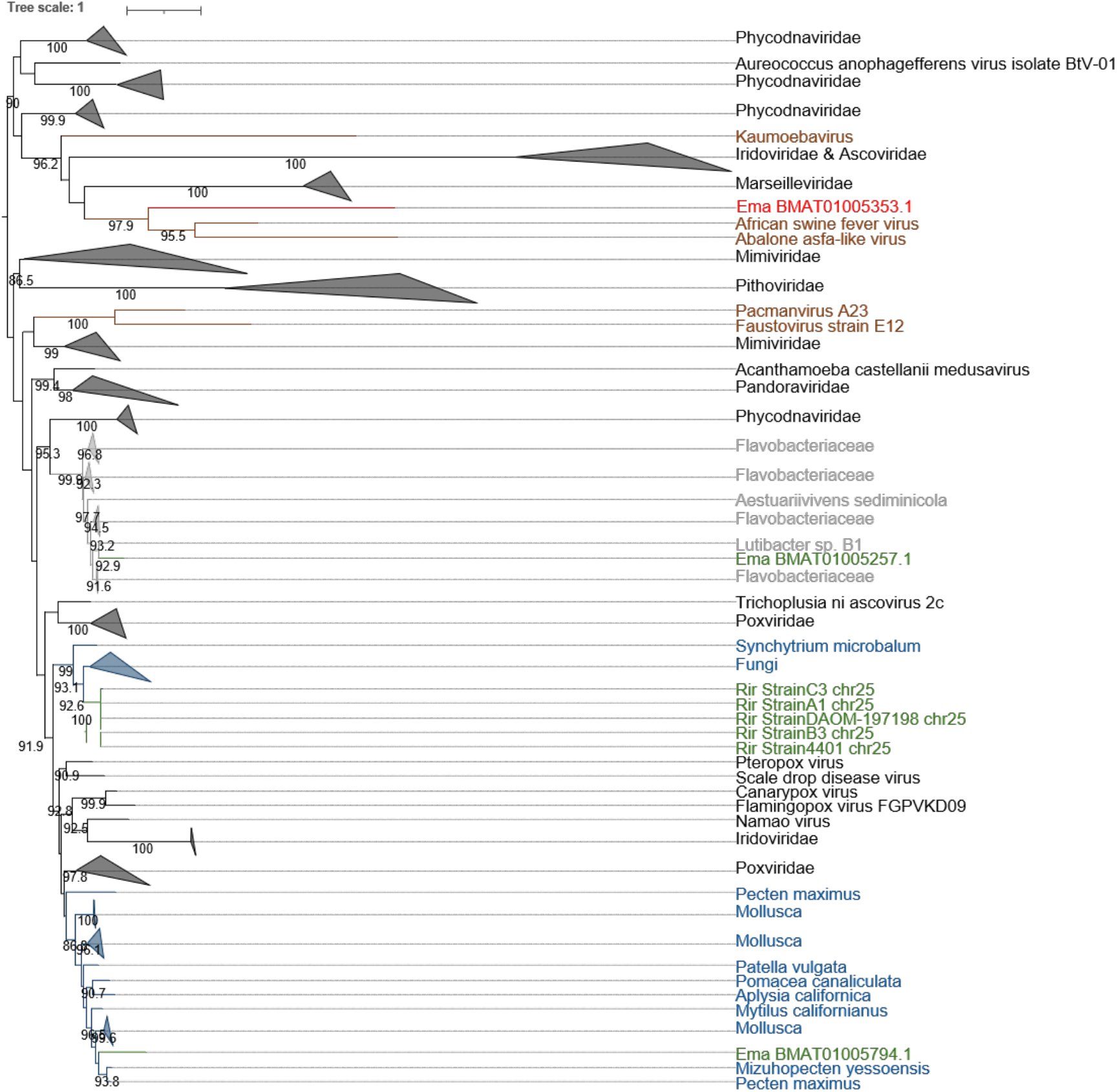
Phylogenetic tree of RNR candidates predicted by ViralRecall. The Fig. S2 legend explains the meaning of the different colors. Ultrafast bootstrap support values are provided along the branches. The root of the tree was arbitrarily chosen and the tree should be considered as an unrooted tree. The best-fit model was Q.yeast+I+I+R7.

**Supplementary Fig S7:**
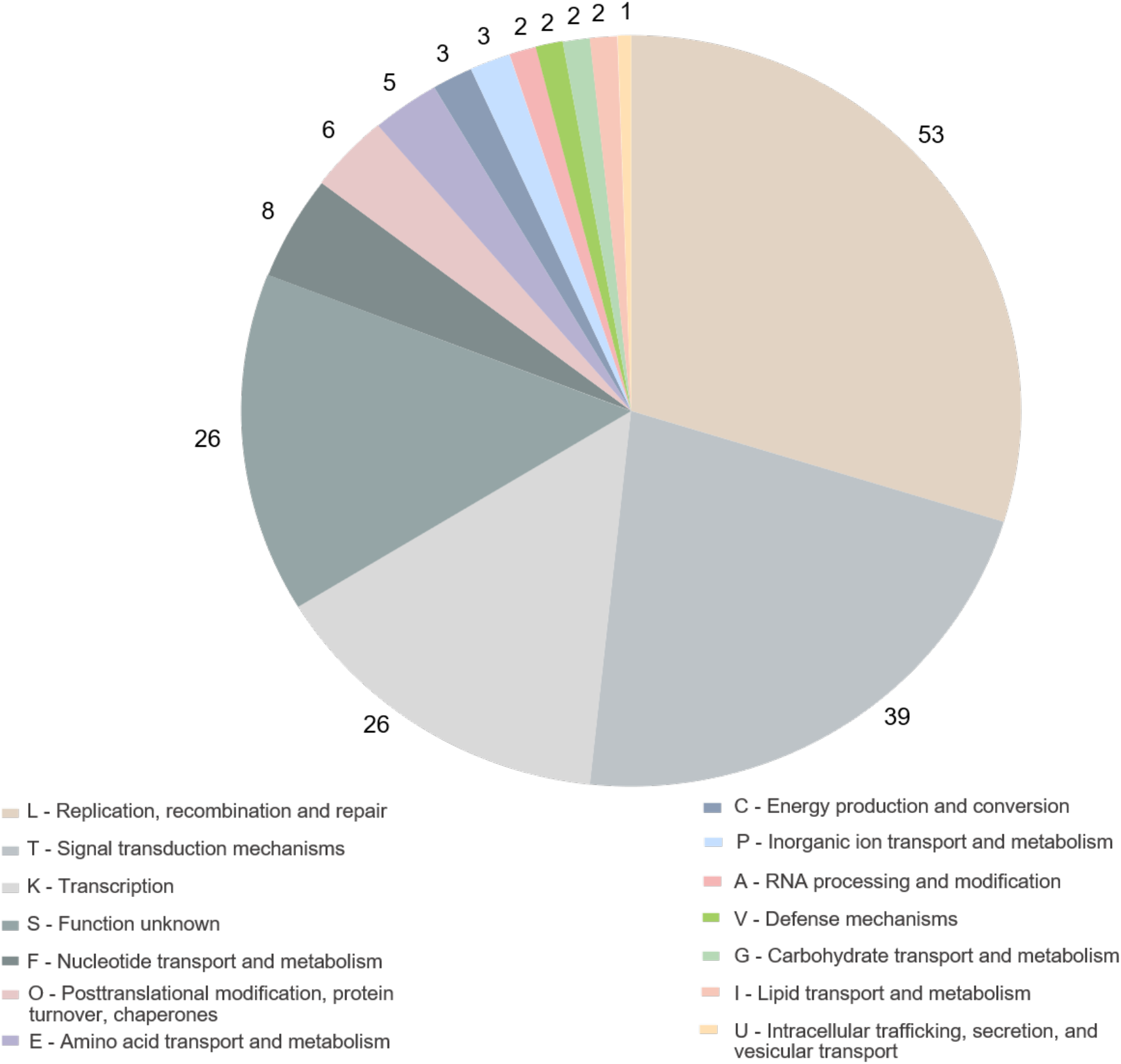
Functional annotation of the GEVE. The number of ORFs belonging to each COG functional category is indicated.

**Supplementary Fig S8:**
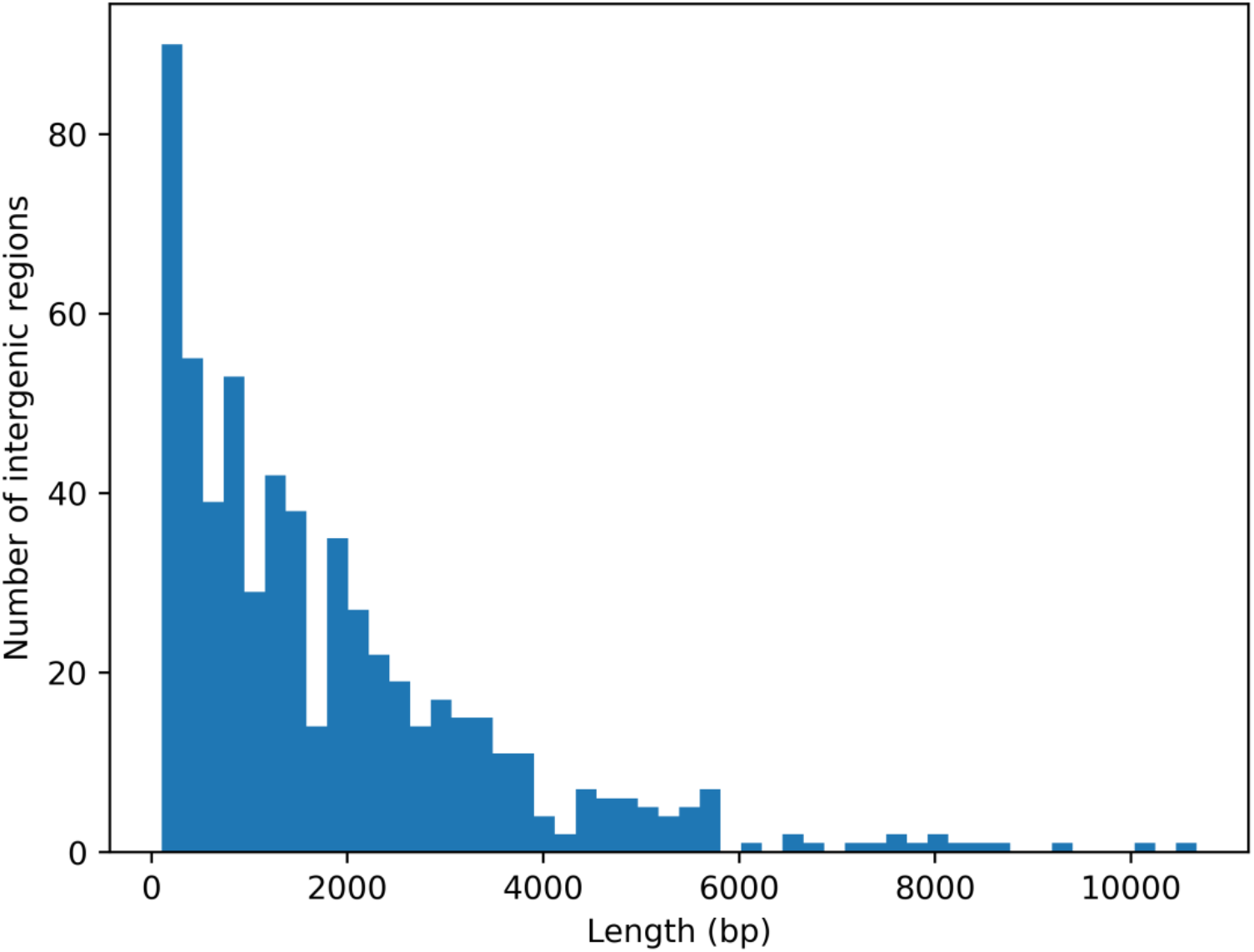
Lengths of the 603 intergenic sequences in the 1.5 Mb GEVE region. The longest intergenic sequence is 10,673 bp. The average length and standard deviation are 1,845 bp and 1,720 bp, respectively.

**Supplementary Fig S9:**
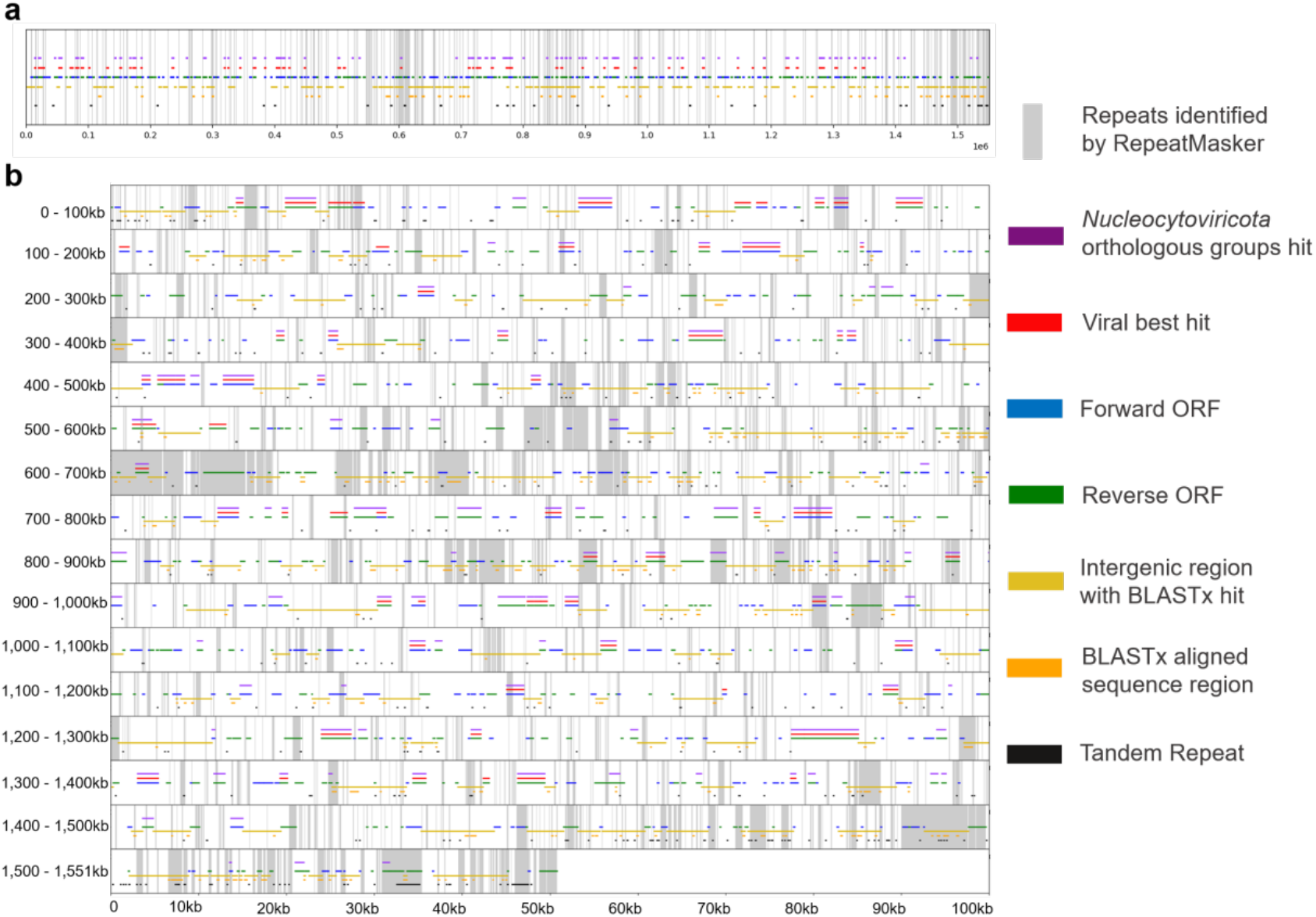
Distribution of genomic components in the 1.5 Mb GEVE region. (a) Overall distribution of genomic components in the GEVE region. Gray vertical lines represent repeats identified by RepeatMasker. Blue and green horizontal lines represent the forward and reverse ORFs predicted by Prodigal, respectively. The red horizontal line represents a sequence that is most similar to a viral sequence (sequences from *E. marginata* were also considered as viral sequences), whereas the purple horizontal line represents a match with *Nucleocytoviricota* orthologous groups in ViralRecall. The yellow horizontal line represents the intergenic region with matched sequences according to the BLASTx results (i.e., traces of genes), whereas the orange horizontal line represents the BLASTx aligned sequence region. The black horizontal line represents the tandem repeat region identified by Tandem Repeat Finder. (b) Details regarding the distribution of genomic components in the 1.5 Mb GEVE region. The colors are the same as in (a). Each row represents 100 kilobases. Some ORFs were identified as repeats by RepeatMasker because of the existence of similar sequences in the fungal genome.

## Supplementary tables

**Supplementary Table S1: Distribution of 10 *Nucleocytoviricota* marker genes predicted by ViralRecall**. The marker gene candidates had HMM scores exceeding the threshold set by ViralRecall (RNAPL, RNAPS, PolB: 200; SFII: 100; Others: 80). The MCP gene was not detected in some viruses, but all five *Asfarviridae* viruses are known to have this gene. These false negatives were probably due to the limitation of the built-in Prodigal and HMM algorithms.

**Supplementary Table S2: Viral regions predicted by ViralRecall in the five *R. irregularis* strains**

**Supplementary Table S3: Distribution of *Nucleocytoviricota* marker genes in the five *R. irregularis* strains**

**Supplementary Table S4: Predicted viral regions and marker genes in the *E. marginata* genome**

